# Deep Learning Strategies for Differential Expansion Microscopy

**DOI:** 10.1101/743682

**Authors:** Domenico L. Gatti, Suzan Arslanturk, Sinan Lal, Bhanu P. Jena

## Abstract

Differential expansion microscopy (DiExM) achieves greater than 500-fold volumetric expansion of biological specimens without loss of cellular antigens. The anisotropic character of this expansion, which affects tissues, cells, organelles and even sub-organelle features, requires the application of novel machine learning approaches to extract accurate and meaningful morphological and biochemical information from DiExM images. Here we describe current strategies to achieve this goal.

---

Expansion Microscopy (ExM) is a novel methodology that uses hydration-competent polymers to expand biological specimens, thus allowing imaging at approximately 70 nanometer resolution using a light microscope^1-3^. The current ExM protocol provides approximately a 64-fold volumetric increase^2^. Although the initial work suggested that ExM is characterized by isotropic expansion, our recent studies show that different tissues expand to different extents and subcellular compartments with different composition undergo anisotropic or differential expansion (DiEx)^4^. In our DiEx microscopy (DiExM) study^4^, greater than 500-fold volumetric expansion is achieved without loss of cellular antigens. DiExM is anisotropic between tissues, organelles, and within organelles themselves. Consequently, there is distortion of the native shape and size of subcellular compartments upon expansion, parameters which are critical in assessing cellular states. Since the relative dimensions and morphologies of cellular components are important parameters commonly used in diagnostic pathology, in DiExM we need to employ both traditional machine learning and deep learning protocols to extract valuable information that may otherwise be lost. We find it to be advantageous to combine applications developed within the Matlab^®^ and Keras/TensorFlow^5,6^ computational platforms, to obtain this information.

For image feature engineering, we have developed a Matlab^®^ Interactive Image Processing Application (MIIPA) that executes sequentially different layers of image processing and analysis (see http://veloce.med.wayne.edu/~gatti/image-processing.html for the complete source code and a video tutorial). Cropping, removal of local artifacts via a hand-drawn mask, and channel selection are user directed. Non-uniform background illumination is corrected by morphological opening^7^. Uncorrelated channel color features (i.e, the blue of DAPI stained nuclei, the green or red of a cellular component decoration) are enhanced by decorrelation stretching (a technique affine to principal component analysis, PCA) based on the eigen decomposition of the channel-to-channel correlation matrix^8^. Additional linear contrast stretching further expands the color range of the resulting image by saturating equal fractions at high and low intensities: typically, the transformed color range is mapped to a normalized interval between 0.01 and 0.99, saturating 2%. Denoising is carried out by passing each channel of an RGB image (or the single luminance channel of an equivalent image in CIE 1976 Lab color space) through a denoising convolutional neural network (DnCNN)^9^ pre-trained on images with added Gaussian noise.

Segmentation of cellular outlines or subcellular components such as nuclei and mitochondria, is possible in MIIPA if a component is specifically decorated in one of the image channels. In this case the channel is binarized by the Otsu method^10^ with a combination of global and adaptive thresholding^11^ with user defined sensitivity, followed by area opening, cycles of dilation, hole filling and erosion of the corresponding mask. In the case of partially overlapping components, visual inspection of the channel is necessary to complement the masks with user defined separating segments and/or hand drawn polygonal mask elements as explained in the <u>Video Tutorial</u>. Upon segmentation, several morphometric parameters are directly extracted from the individual components including area, bounding box, centroid, pixel values with maximal, minimal, mean and s.d. intensity, weighted centroid based on pixels intensity, circularity, eccentricity, equivalent diameter, major and minor axis length, orientation, perimeter and solidity.

A unique MIIPA feature is the utilization of Patterson maps to determine the radial distribution of one subcellular component (i.e., mitochondria) around another one (i.e., nuclei), if a segmentation mask for these components has been calculated. The Patterson map^12^ of an image is obtained by calculating the fast fourier transform (FFT) of the image, and then back-transforming (*via* inverse FFT, iFFT) removing the phases and using only the squared amplitudes. The map contains all the vectors from every pixel of the image to every other pixel, weighted by the pixel intensity. Thus, for example, subtracting from the Patterson map of the nuclei + mitochondria (containing intra-nuclei, nuclei-to-nuclei, nuclei-to-mitochondria, intra-mitochondria, and mitochondria-to-mitochondria vectors) the Patterson map of the nuclei (containing only intra-nuclei and nuclei-to-nuclei vectors) and the Patterson map of the mitochondria (containing only intra-mitochondria and mitochondria-to-mitochondria vectors) leaves only the nuclei-to-mitochondria vectors. The resulting map is called a Difference Patterson map (**Figure 1**). Regions of this map with high vector density reflect a strong mitochondrial signal. Radial averaging of the map provides the distribution of mitochondria around the nuclei centroids.

**Figure 1.**
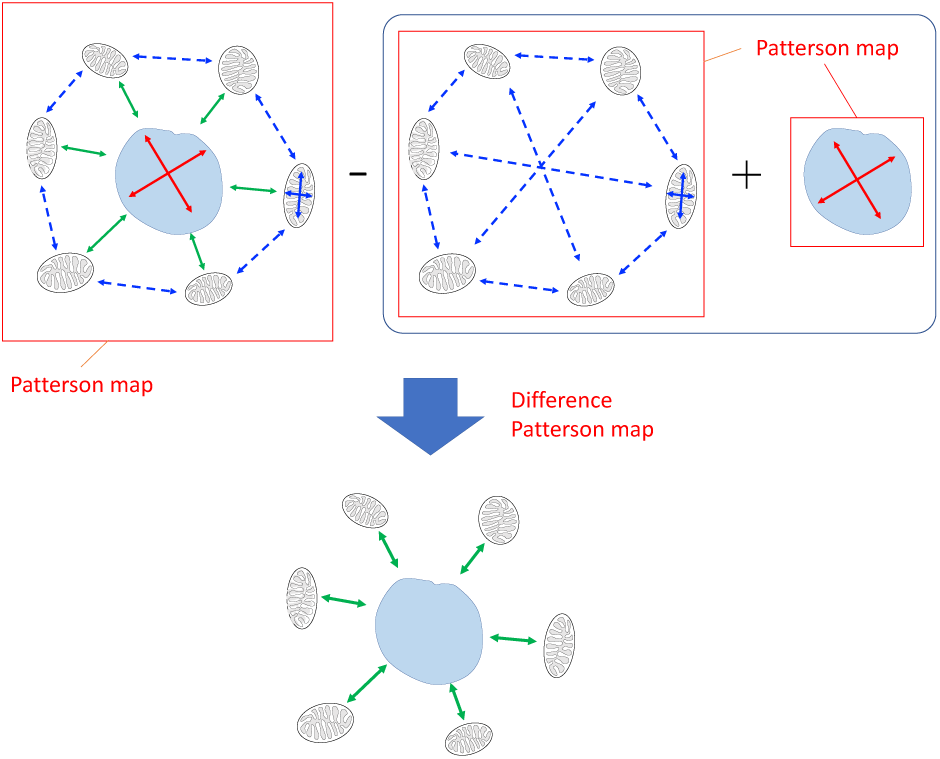
Radial distribution of mitochondria around a nucleus. The sum of the Patterson map of the mitochondria (containing only intra-mitochondria and mitochondria-to-mitochondria vectors) and the Patterson map of the nucleus (containing only intra-nucleus vectors) is subtracted from the Patterson map of the nucleus + mitochondria (containing intra nucleus, nucleus-to-mitochondria, intra-mitochondria, and mitochondria-to-mitochondria vectors), leaving only the mitochondria-to-nucleus vectors.

In a similar way, MIIPA uses Patterson maps to calculate density functions for subcellular components. For example, radial average of a difference Patterson map calculated by subtracting from the Patterson map of all nuclei (containing all vectors within and between all nuclei) the sum of the Patterson map of an individual nucleus (all vectors inside that nucleus) and the Patterson map of all nuclei except that nucleus provides the distribution of the distances of that nucleus from every other nucleus. Final averaging of the Difference Patterson maps for all nuclei in an image provides the distribution of the inter-nuclei distances. These calculations make use of the FFT and iFFT, and thus are extremely fast.

Numerical information about the cells and their subcellular compartments derived by MIIPA is then processed via traditional supervised (i.e., classification) or unsupervised (i.e., PCA) machine learning applications or fed to a separate input port of an artificial Neural Network designed to analyze DiExM images. Although Deep Learning protocol based on Convolutional Neural Networks (CNN) are readily developed and/or built using Matlab^®^ Deep Learning Toolbox, because some NN architectures are not supported yet by this Toolbox, we currently prefer to develop Deep Learning applications for DiExM using the Keras/Tensorflow platform^5,6^. An example of a NN architecture used for classification of DiExM images is shown in **Figure 2**. Images of 1040 × 1388 pixels are preprocessed to size 260 × 347 pixels. The left branch of the network consists of a convolutional sub-network^13^ (CNN) that processes these images, reducing each image to a vector of 128 scalar values. The aforementioned MIIPA derived numerical data associated with each image is in the form of a vector of 805 scalar values containing information on the distributions of nuclei areas, long and short axes, the distribution of inter-nuclei distances, the mean cell area, and the mean nucleus to cell area ratio. The right branch of the network consist of a dense sub-network^14^ (DNN) that processes this numerical data and reduces it to a vector of 64 scalar values. Concatenation of the CNN and DNN output vectors results in a vector of 192 scalar values, which by application of a *softmax* function leads to the assignment of each image to one of several classes (**Figure 2**). We are currently training this NN to classify DiExM images according not only to normal/diseased states of the biological samples, but also to specific features of the expansion protocol, like the percentage of fixative used and the level of expansion achieved.

**Figure 2.**
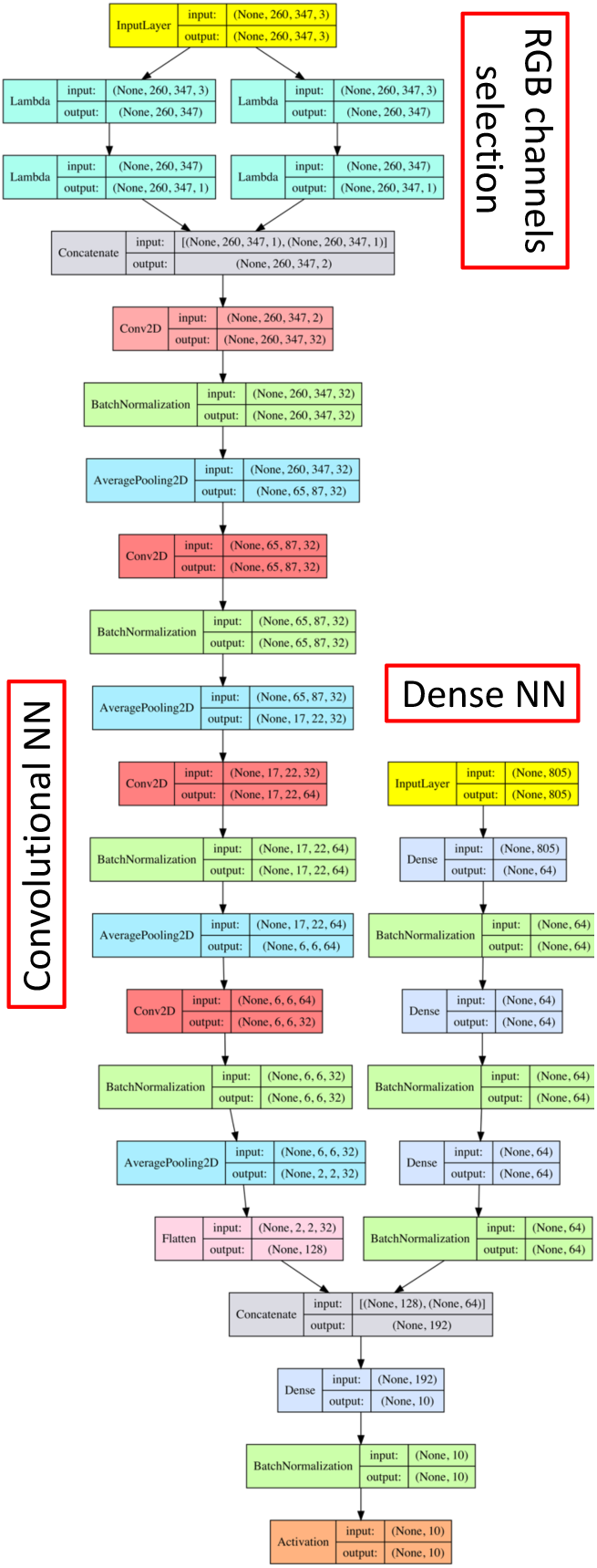
Architecture of the mixed-type input NN used to classify DiExM images. The left branch of the network is a *convolutional* sub-network (CNN) that reduces each image to a vector of 128 scalar values. The right branch of the network is a *dense* sub-network (DNN) that reduces image associated numerical data to a vector of 64 scalar values. Concatenation of the CNN and DNN outputs results in a vector of 192 scalar values that progresses in a common *dense* path ending with a *softmax* layer for the assignment of each image, and the associated numerical data to one of several classes.

With modification only in the input and final output, the same architecture can be used for *regression* instead of *classification*. For example, at this time, due to DiExM image to image variation depending on various factors (i.e., resolution, brightness, type of staining, number of channels, cell density, etc.), Matlab® based MIIPA still requires user intervention for fine-tuning the segmentation of organelles and for the adjustments of various parameters used in image analysis. While MIIPA is robust enough that the ranges of parameter values for optimal performance are quite large, we are currently working toward completely automating the feature engineering process. For this purpose, a separate CNN is being trained with DiExM images that have been hand-labeled with the correct set of parameters for optimal feature extraction. In this case, the CNN will work as a *regressor*, rather than a *classifier*, providing as the output of its final layer the required set of parameters for input to MIIPA. The anticipated pipeline for automated DiExM image analysis is shown in **Figure 3**.

**Figure 3.**
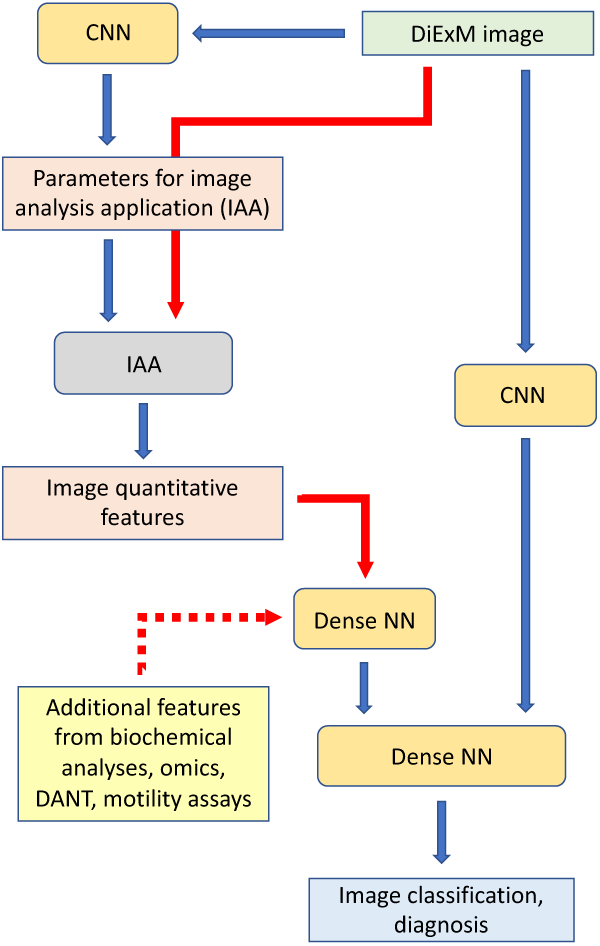
Pipeline for diagnostic assignment of DiExM images. DiExM images are first passed through a CNN to assign parameters required by an image analysis application (IAA) that produces a vector of quantitative features. These features (as well as additional features from other potential sources) are then channeled through a dense network and finally merged with a separate CNN that detects morphological characteristics. The output of the pipeline is a diagnostic class assignment.

An alternative version of this pipeline transfers all semantic segmentation operations from the image analysis application (i.e., our MIIPA) to a dedicated NN. For this task we adopt the architecture of Res-U-Net^15^, a type of *fully convolutional* NN^16^ designed for road area extraction. This architecture combines the low level detail information and high level semantic information achieved by U-Net^17^, a type of *encoder-decoder* NN^18^, which has shown high performance in biomedical image segmentation, with the ease of training and resilience to gradient degradation of *residual* neural networks^19^. A residual network consists of a series of stacked residual units, each unit being represented by the general expression:

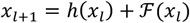

where *x*_*l*_ and *x*_*l*+1_ are the input and output of the *𝓁* -th residual unit, ℱ is the residual function, and *h*(*x*_*l*_) is an identity mapping function (i.e., *h*(*x*_*l*_) = *x*_*l*_). The flowchart of a typical residual unit in Res-U-Net is shown in **Figure 4.**

**Figure 4.**
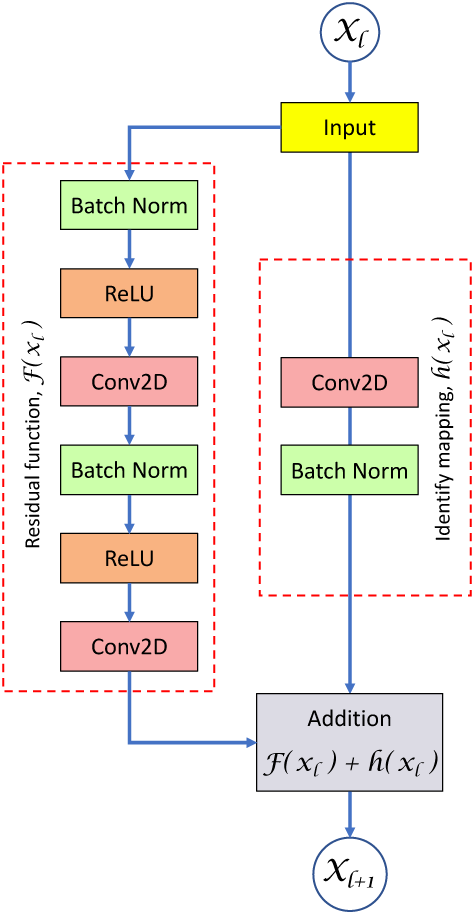
The residual unit with identity mapping used in Res-U-Net. The left path transforms the input feature map through a series of convolutions and nonlinear activations extracting semantic information. The right path, often called a *skip connection*, simply passes the feature map through without any large transformation except those (i.e., convolutions, batch normalizations) necessary to match the size of the output from the left path. The paths are then added together, which allows the network to learn subtle transformations (the residual function), or if necessary to reduce the training error by driving the weights of the left path toward zero and leaving only the identity mapping.

In Res-U-net, the skip connections within a residual unit and between low levels and high levels of the network facilitate the propagation of information without degradation, making it possible to design a neural network with fewer parameters than U-Net, but with better performance on semantic segmentation. The network comprises of three parts: *encoding, bridge*, and *decoding*. The first part encodes the input image into compact representations conceptually similar to the words *embeddings* used in natural language processing (NLP)^20^. The last part recovers the representations to a pixel-wise categorization, i.e. a semantic segmentation. The middle part acts like a bridge connecting the encoding and decoding paths. All of the three parts are built with residual units as shown in **Figure 4**. The encoding path has three residual units. In each unit, instead of using pooling operation to down-sample the feature map size, a stride of 2 is applied to the first convolution block to reduce the feature map by half. The decoding path consists also of three residual units. Before each unit, there is an up-sampling of feature maps from lower level and a concatenation with the feature maps from the corresponding encoding path. After the last level of decoding path, a 1×1 convolution and a *sigmoid* or *softmax* activation layer are used to project the feature maps into the desired segmentation. As the loss function, we use a combination of the cross entropy loss and of the Dice coefficient^21,22^, which has been shown to increase performance over just the cross entropy loss. The Dice coefficient can be generalized to continuous binary vectors either by the summation of the probabilities in the denominator or by the summation of their squared values as shown below:

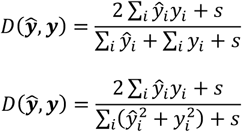

Herein, ***Ŷ*** ≡ {*Ŷ*_*i*_}, *Ŷ*_*i*_ ∈ [0,1] is a continuous variable, representing the vector of probabilities for the *i*-th pixel, ***y*** ≡ {*y*_*i*_}, are the corresponding ground truth labels, and *s* is a *Laplacian* smoothing scalar. For binary vectors (i.e., segmentation masks used as labels), *y*_*i*_ ∈ {0,1}. The combined binary cross-entropy + Dice loss is thus derived as:

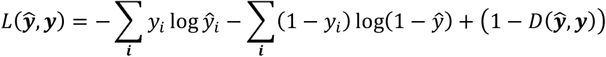

A flowchart of the Res-U-Net used for segmentation of DiExM images is show in **Figure 5**. Training of this network uses as labels segmentation masks for either cells or subcellular components produced by an initial MIIPA run on the same DiExM images.

**Figure 5.**
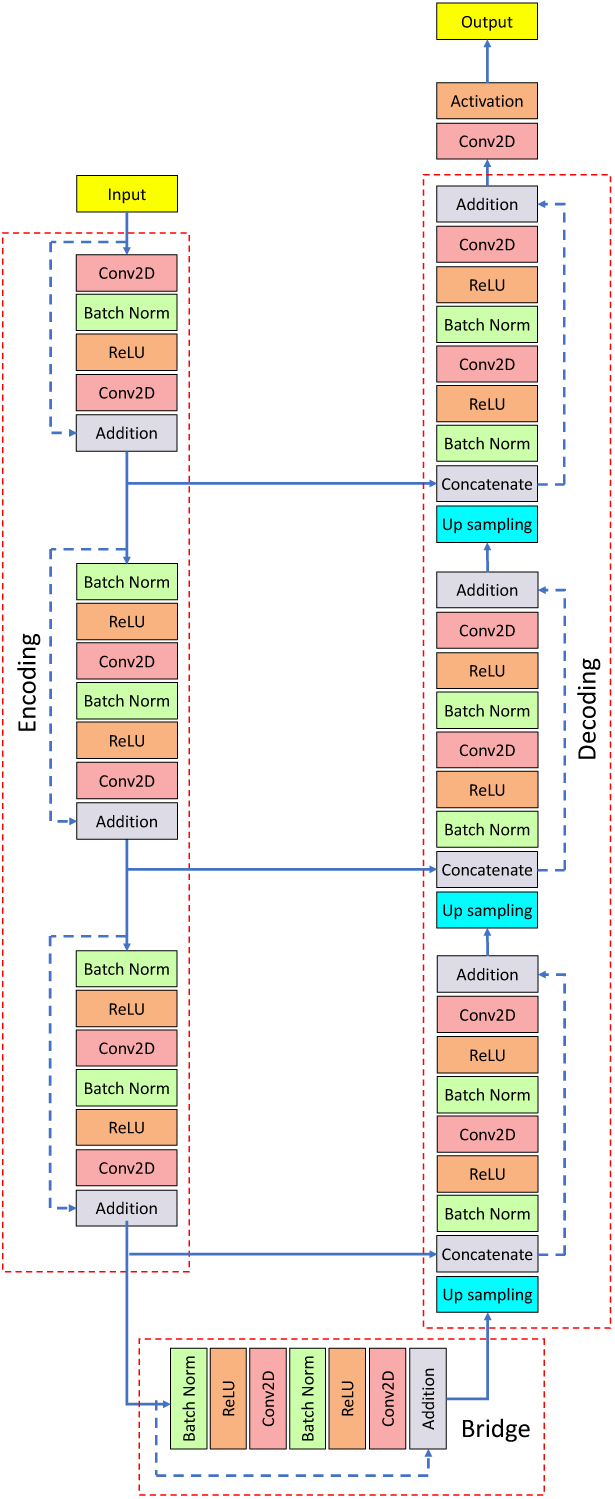
Flowchart of a Res-U-Net fully convolutional neural network for semantic segmentation. A U-Net (so called for its U-shaped architecture) is a classic way to arrange operations for segmenting and denoising images. In Res-U-Net, several residual blocks (Figure 4) are arranged in sequence. In each block of the left-hand side of the network the feature map is down-sampled. After three residual blocks, the resulting map containing the encoded semantic information is passed to the right-hand side of the network through a *bridge* res-block and iteratively up-sampled to bring the resolution to that of the original image. In order to increase localization (the assignment of a class label to each pixel), high resolution features from the *encoding/contracting* path are concatenated with the up-sampled features of the *decoding/expansive* path through a skip-connection before being finally convolved and output into a binary mask.

Another application of a *regression* oriented CNN that we are actively developing is the extraction of *biochemical information* from DiExM images of skeletal muscle cells in culture. A widely accepted paradigm of cell biology is that cell morphology and function are not only ‘interconnected’, but mutually ‘determining’. In no other cells is this connection more evident than in muscle cells, where the spatial localization of energy producing mitochondria near myosin fibers and axonal ending is critical for contraction and communication. On this basis, we are using *Deep Learning* to reveal the correlation between the metabolic activity of muscle cells and their DiExM images. Skeletal muscles account for more than a third of our body weight, and myocytes metabolism has a major impact on whole-body homeostasis. For instance, myocytes are responsible for ∼75% of the insulin-stimulated clearance of glucose from the blood after a meal^**23**^. Mitochondria are key determinants of muscle cell functions controlling the spatiotemporal supply of energy. In this context, using human myocytes differentiated into myotubes in culture we can assess accurately the mitochondrial respiratory activity by high resolution oximetry and the fluxes of major reporter metabolites for glycolysis, pentose shunt, and TCA cycle using SeaHorse technology (Agilent Technologies, Inc.) and other traditional biochemical/spectroscopic assays. Using a small number of experimentally derived parameters as constraints, we can then derive computationally a map of the entire myocyte metabolism using the recently reconstructed genome-scale metabolic model (GEM) of human myocytes, iMyocyte2419^24^, which includes 5,590 reactions and 4,448 metabolites in 8 different compartments. This reconstruction exploits the theoretical frame of Flux Balance Analysis (FBA)^25^. In FBA the traditional representation of the vector of time derivatives of metabolites concentrations ***x*** as the product of a *rate matrix* ***K*** and ***x*** is replaced by the product of a *stoichiometric matrix* ***S*** and a vector ***v*** of *fluxes*:

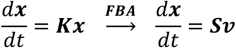

The *stoichiometric matrix* is organized such that every column corresponds to a reaction and every row corresponds to a compound, and thus it contains information on all the chemical reactions taking place in a cell. The entries in the matrix are *stoichiometric coefficients*, which are integers. Every row describes the reactions in which that compound participates and therefore how the reactions are connected by it. Therefore, the *dynamic mass balance equation*:

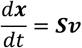

describes all possible *dynamic* states of the metabolic system, with the *flux balance equation*:

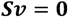

representing all possible combinations of metabolic fluxes consistent with a *steady-state*. Flux Balance Analysis seeks to maximize or minimize an objective (cost) function:

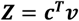

where ***c*** is a vector of weights indicating how much each reaction contributes to the objective function. In practice, when only one reaction is desired for maximization or minimization, ***c*** is a vector of 0 with a value of 1 at the position of the reaction of interest. Optimization of such a system is accomplished by a class of algorithms known as *linear programming*^*26*^. FBA can thus be defined as the use of linear programming to identify the best cost function:

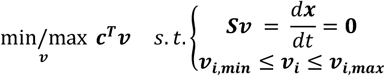

where ***v*** _***i***, ***min***_ ≤ ***v*** _***i***_ ≤ ***v*** _***i***, ***max***_ are upper and lower limits on particular fluxes of metabolites as determined experimentally (see above). The output of FBA is a particular flux distribution, ***v***, which maximizes or minimizes the objective function. Thus, the most important step in the application of FBA is the choice of the *objective function* that allows the identification of a particular functional state in the space of possible solutions. For example, if the cultured myotubes are in log-phase of growth we use as objective function the maximization of the biomass synthetic reaction to reconstruct the entire metabolism. If the cultured myotubes are in stationary phase we reconstruct their metabolism using as objective function the minimization of energy demand and therefore ATP synthesis.

FBA metabolic reconstruction of myocytes in culture is the key preliminary step in the development of our ambitious goal and completely novel approach to understand the relationship between *form* and *function* in these cells. On this basis, we use a *regression output* variation of our mixed-type input NN to reveal the correlation between the metabolic activity of muscle cells and their DiExM images. The vectors of biochemical *markers* of metabolic fluxes collected from cultured muscle cells are used as the labels of the corresponding DiExM images. Our NN is then trained to associate a particular metabolic vector to a specific DiExM image. When a previously unseen DiExM image is passed through the NN, the regression output provides the vector of marker metabolic fluxes that can be used as constraints to reconstruct the metabolic state of the myocytes simply from their DiExM images, without the need for all the biochemical analyses that were necessary to initially train the network. In the future, this approach will be used to infer the metabolic state of cells from different tissues in diseased individuals simply from imaged biopsy material, without the need for laborious, expensive or invasive biochemical tests.

Finally, we stress that in order to avoid overfitting and increase sample size, we regularly apply geometric image *augmentation* and *shuffling* in the training of our NN’s for classification, segmentation, and regression of DiExM images so that, in each iteration, the NN never sees the exact same set of images. Random augmentation modes include shifting, down/up-sizing, rotating, shearing, and mirroring operations. The parts of an image that are left out from the frame after the transformation are filled in with 0’s. For segmentation, each pair of image and ground truth mask is modified the same way. For regression and classification, augmented images retain the same labels.

DiExM is a novel methodology that promises to revolutionize clinical pathology by achieving nanometer scale resolution with diffraction limited optical microscopy. The full investigative potential of DiExM is reached when morphological and biochemical information of cellular structure/function and diagnostic significance is comprehensively extracted from DiExM images with both protocols of traditional Machine Learning and various architectures of Deep Learning neural networks.

## Author Contributions

D.L.G., B.P.J. and S.A. developed the idea. D.L.G. wrote the manuscript. S.L. participated in the study. All authors participated in discussions and proofreading the manuscript.

## Acknowledgements

Work presented in this article was supported in part by the National Science Foundation grants EB00303, CBET1066661 (BPJ).

